# Microphysiological system modeling pericyte-induced temozolomide resistance in glioblastoma

**DOI:** 10.1101/2024.07.16.603611

**Authors:** Surjendu Maity, Christopher Jewell, Can Yilgor, Satoru Kawakita, Saurabh Sharma, Alejandro Gomez, Marvin Mecwan, Natashya Falcone, Menekse Ermis, Mahsa Monirizad, Negar Hosseinzadeh Kouchehbaghi, Fatemeh Zehtabi, Danial Khorsandi, Mehmet Remzi Dokmeci, Diogo Moniz-Garcia, Alfredo Quiñones-Hinojosa, Ali Khademhosseini, Vadim Jucaud

## Abstract

Glioblastoma (GBM) is a malignancy with poor survival and high rates of chemoresistance. Temozolomide (TMZ), the standard-of-care chemotherapy for GBM patients, but GBM cells can be resistant to TMZ, resulting in limited clinical efficacy. Elucidating the complex mechanisms of TMZ chemoresistance in GBM requires novel *in vitro* models replicating the complex tumor microenvironment (TME). We present an multicellular 3D GBM model recapitulating the biomechanical characteristics of brain tissues and pericyte-mediated TMZ resistance. The composite hydrogel used to encapsulate GBM spheroids (U87, LN229, and PDM140), pericytes, or GBM spheroids with pericytes, mimics the rheological properties of brain tissues (G’∼800Pa and G”∼100Pa). When untreated, the GBM models remain viable and proliferative for 14 days. PDM140 spheroids were most sensitive to TMZ (IC_50_=73μM), followed by LN229 (IC_50_=278μM) and U87 (IC_50_=446μM). With pericytes, the viability of TMZ-treated GBM spheroids significantly increases by 22.7% for PDM140, 32.5% for LN229, and 22.1% for U87, confirming pericyte-induced GBM chemoresistance responses. The upregulation (380-fold) of C-C motif chemokine ligand 5 (CCL5) in pericytes upon TMZ treatment could explain the chemoresistance responses. This innovative brain-mimicking 3D GBM model represents a novel *in vitro* platform for testing the efficacy of TMZ and novel drugs targeting CCL5-mediated chemoresistance pathways in GBM.

## 1. Introduction

Glioblastoma multiforme (GBM) is a grade IV glioma that represents approximately 30% of all primary brain tumors. GBM is the most common and malignant type of glioma, with an annual incidence of 5.26 cases per 100,000 people associated with a high economic burden accounting for about 13,000 deaths per year and $7.2 billion in spending in the U.S. alone.^[1]^ With the current standard-of-care treatment for GBM, including maximal tumor resection followed by concomitant radiotherapy and temozolomide (TMZ) chemotherapy, the median survival is 14.6 months, with a 2-year survival of 27% and a strikingly poor 5-year survival of 4.7%.^[1b-d]^ TMZ is a blood-brain barrier (BBB)-penetrating DNA alkylating agent that attaches a methyl group at the O^6^, N^7^, and N^3^ position of guanine residues, which causes DNA damage and cell-cycle arrest in the G2/M phase to induce the apoptosis of GBM cells.^[2]^ However, the innate and adaptive capacity of GBM cells to be resistant to TMZ results in limited clinical efficacy, as manifested by >50% of GBM patients not responding to the treatment.^[3]^ Consequently, GBM chemoresistance remains a major clinical obstacle for successfully managing GBM patients with currently available therapies, contributing to treatment failure and disease recurrence.^[4]^

The biological basis of TMZ resistance has been an active research area, and several mechanisms have already been uncovered. However, minimal progress toward better treatments has been achieved as all the mechanisms of TMZ resistance have not been fully elucidated, and further investigations are required to identify novel therapeutic targets and treatment strategies.^[5]^ Resistance to TMZ can arise through various mechanisms reflecting the heterogeneity and adaptability of cancer cells within the tumor microenvironment (TME). The primary mechanisms contributing to TMZ resistance in GBM are enhanced DNA repair mechanisms where TMZ-resistant GBM cells upregulate proteins involved in the mismatch repair pathway,^[6]^ such as MGMT (O6-methylguanine-DNA methyltransferase), and the DNA base excision repair pathway,^[7]^ which lessen the cytotoxic effects of TMZ and promote tumor cell survival. Other mechanisms contributing to TMZ resistance in GBM include alterations in drug metabolism, dysregulation of apoptotic pathways, and pro-survival signaling cascade activation.^[2b, 4]^ Lastly, the heterogeneous nature of GBM tumors, with high levels of inter- and intra-tumor heterogeneity, contributes to the complexity of TMZ resistance, as different subpopulations of GBM cells (i.e., proneural, classical, and mesenchymal) may exhibit distinct molecular profiles and responses to therapy.^[8]^

In addition to genetic and epigenetic alterations within GBM cells, the GBM TME is a highly heterogeneous and complex system composed of different cell populations and extracellular matrix (ECM) components that play a crucial role in GBM progression and treatment response.^[8-9]^ Besides GBM cells, which are the primary drivers of tumor growth and invasion, the GBM TME includes stromal cells (i.e., pericytes, endothelial cells, and fibroblasts) that contribute to angiogenesis, extracellular matrix remodeling, and immune modulation within the TME,^[10]^ immune cells (i.e., macrophages, microglia, and lymphocytes) that play a role in promoting or suppressing tumor growth,^[11]^ and astrocytes and neural precursor cells that influence tumor progression and therapeutic resistance.^[12]^ The GBM TME also includes an ECM composed of a complex macromolecule network distinct from healthy brain tissues.^[13]^ Indeed, there is an increased expression of specific ECM components in the GBM TME (i.e., hyaluronic acid (HA), glycosaminoglycans, thrombospondin, and fibronectin), leading to changes in drug penetration (e.g., TMZ), immune cell recruiting and modulation, and signaling pathway activation, that contribute to tumor invasiveness and development.^[14]^ Therefore, elucidating the complex mechanisms of TMZ resistance driven by the GBM TME is essential for developing targeted therapies to overcome treatment failure and improve GBM patient outcomes.

Within the GBM TME, pericytes are multipotent perivascular cells essential for tumor initiation, survival, and progression.^[15]^ Pericytes are integral to the vascular GBM niche by promoting vascular stability, integrity, and function. Alterations in pericyte coverage, distribution, phenotype, and function have been observed in GBM, contributing to aberrant angiogenesis, vascular dysfunction, and treatment resistance.^[16]^ In addition, pericytes can promote the growth of cancer cells and drug resistance through paracrine signaling.^[17]^ In particular, pericytes can interact with GBM cells to enhance DNA repair and induce TMZ resistance through the C-C motif chemokine ligand 5 (CCL5)-C-C motif chemokine receptor 5 (CCR5) paracrine signaling pathway, where pericytes secrete CCL5 that subsequently binds to the CCR5 expressed by GBM cells and activate the DNA damage response upon TMZ treatment.^[18]^ Although the function of the CCL5-CCR5 paracrine signaling in TMZ resistance remains to be fully elucidated, the excessive outgrowth of pericytes and the overexpression of CCL5 in GBM patients highlights the potential of the CCL5-CCR5 paracrine axis as a critical driver of TMZ resistance.^[19]^ Therefore, understanding the crosstalk between pericytes and GBM cells, particularly in TMZ resistance, may uncover novel therapeutic target opportunities to enhance treatment efficacy and improve GBM patient outcomes.^[18, 20]^

A critical factor that impedes progress toward GBM drug discovery, development, and screening is the paucity of simple and reliable models that recapitulate the complex mechanisms of TMZ resistance driven by the GBM TME,^[21]^ particularly pericyte-induced TMZ resistance in GBM. Current two-dimensional (2D) cell culture models do not recapitulate the complex physiology of the GBM TME,^[22]^ and preclinical animal models can lead to inaccurate predictions of human GBM biology and drug responses.^[23]^ To address these limitations, 3D microphysiological systems (MPS) have emerged as powerful tools for studying complex biological processes and disease mechanisms *in vitro*.^[24]^ These innovative platforms enable the recreation of physiologically relevant microenvironments by integrating multiple cell types, including tumor cells, stromal cells, vascular cells, and ECM components. Current *in vitro* 3D GBM MPS can mimic tumor growth, migration, and therapy response, thus offering biologically relevant platforms recreating *in vivo* physiology with accurate spatial cell arrangements, cell-to-cell and cell-to-ECM interactions, and biochemical gradients.^[25]^ However, the physicochemical properties of GBM-specific tissues are rarely considered when developing 3D GBM MPS despite their essential role during neurodevelopment and homeostasis.^[26]^

Hydrogels composed of gelatins, HA, collagen, and their derivatives, including gelatin methacryloyl (GelMA) and hyaluronic acid methacrylate (HAMA), have been used to develop 3D GBM models for their mechanical properties and cellular functionality that replicate the native GBM TME.^[27]^ These hydrogels allow the development of intricate *in vitro* models with biologically relevant cellular dynamics, signaling cascades, and drug responses.^[28]^ Therefore, we developed a multicellular 3D GBM MPS mimicking the physicochemical properties of brain tissues and pericyte-mediated TMZ resistance. We used GelMA and HAMA to develop a biomimetic hydrogel matrix with the rheological properties of brain tissues for encapsulating GBM spheroids and pericytes. Human GBM cells (LN229, PDM140, and U87) and primary pericytes were used as model cell types to demonstrate TMZ resistance heterogeneity between GBM cell lines and healthy cells. We generated different GelMA/HAMA-based 3D bioconstructs containing GBM spheroids, pericytes, or GBM spheroids co-cultured with pericytes. After optimizing the 3D GBM models to remain viable and proliferative for 14 days, we analyzed the viability and chemoresistant-related gene expression of the bioconstructs in response to a 7-day dose-dependent TMZ treatment. Lastly, we compared the cell viability of GBM spheroids cultured with and without pericytes to analyze and confirm pericyte-induced GBM chemoresistance responses *in vitro*.

## 2. Results

### 2.1. Characterization of GelMA-HAMA hydrogels mimicking the rheological properties of brain tissues

To develop a hydrogel matrix with the rheological properties of brain tissues (300<G’<1200Pa and 20<G”<120Pa),^[29]^ we used GelMA and HAMA to develop a composite hydrogel for encapsulating GBM spheroids and pericytes (**Figure 1A**). We synthesized four different composite hydrogels dissolved in cell culture medium with 1% Irgacure: 5% GelMA / 2% HAMA (5G/2H); 3% GelMA / 1% HAMA (3G/1H); 2.5% GelMA / 2.5% HAMA (2.5G/2.5H); and 2% GelMA / 1% HAMA (2G/1H). We confirmed using NMR and FTIR that both GelMA and HAMA biopolymers were methacrylated (**Figure S2A-C**). Subsequently, we generated large cylindrical GelMA/HAMA hydrogel constructs (diameter: 5 mm; and height: 5 mm) with a volume of 98 μL. The rheological analysis of the different hydrogel formulations, crosslinked with UV for 15 seconds, demonstrated that the hydrogel 5G/2H had the highest storage and loss modulus (G’∼2000 Pa and G” ∼200 Pa, respectively), followed by 2.5G/2.5H (G’∼1500 Pa and G” ∼100 Pa), 3G/1H (G’∼800 Pa and G” ∼100 Pa), and 2G/1H (G’∼180 Pa and G” ∼100 Pa) (**Figure 1B and 1C**). The storage and loss modulus of the 3G/1H hydrogel was within the range of brain tissues (300<G’<1200Pa and 20<G”<120Pa);^[29]^ therefore, this hydrogel formulation was used for additional rheological characterization using temperature, frequency, and amplitude sweeps with varying UV crosslinking times (5, 15, 20, 30, or 60 seconds). The storage and loss modulus of the 3G/1H hydrogel was relatively stable between 22°C and 45°C (**Figures 1D and 1E**). At 37°C, the frequency sweep revealed that the storage and loss modulus increased with longer UV crosslinking times: G’∼100 Pa and G” ∼20 Pa after 5 s; G’∼800 Pa and G” ∼100 Pa after 15 s; G’∼2000 Pa and G” ∼400 Pa after 20 s; and after 30 s of crosslinking, the storage and loss modulus plateaued (G’∼5000 Pa and G” ∼1000 Pa) (**Figure 1F and 1G**). Similarly, the amplitude sweep showed that the storage and loss modulus increased with UV crosslinking times from 5 s to 20 s (**Figure 1H and 1I**). The swelling and degradation rates were evaluated for 48 hours and 7 weeks, respectively. Within the first 3 hours, the 3G/1H hydrogel reached a 60% swelling ratio, which plateaued at 80% after 20 hours (**Figure 1J**). The hydrogel degraded at a rate of 0.68 %/day in the first 4 weeks and 0.17 %/day from 4 to 7 weeks to reach a final degradation of 23% after 7 weeks (**Figure 1K**). We also analyzed the effect of crosslinking time on porosity (**Figure 1L**). We observed that the porosity decreased with crosslinking times, with a porosity of 0.38 after 5 s, 0.25 after 15 s, and 0.19 after 20 s (**Figure 1M**). Lastly, a simulation was performed to evaluate the diffusion of TMZ molecules (194.151 g/mol) in the 3G/H1 hydrogel. At all concentrations tested (1, 10, 50, 100, 250, 500, and 1000 μM), TMZ molecules completely diffused through the 3G/H1 hydrogel in 26 hours, respectively (**Figure S1E**). The simulation results were confirmed experimentally using a 120 μM Rhodamine solution (**Figure S1H**). Overall, we confirmed that the 3G/1H hydrogel, crosslinked for 15 s, retains the rheological properties of brain tissues at 37°C (body temperature), displays a porosity that allows the diffusion of small molecules, and slowly degrades over time. Therefore, the 3G/H1 hydrogel formulation was used as the primary matrix to encapsulate GBM spheroids and pericytes in subsequent experiments.

**Figure 1:**
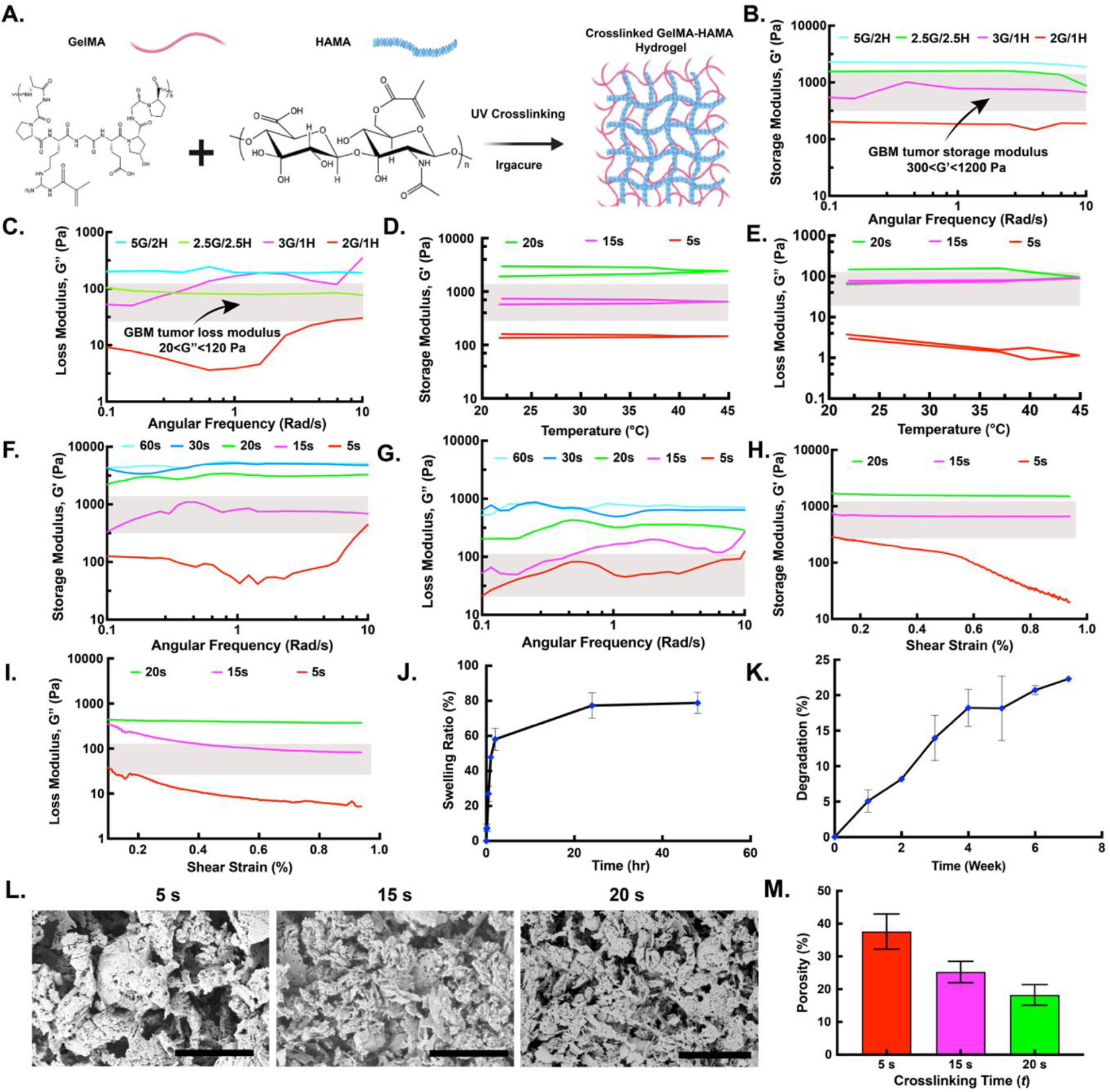
Optimization and characterization of GBM/brain tissue mimicking GelMA-HAMA hydrogel. **A**. Synthesis of GelMA-HAMA composite hydrogels. Frequency sweep analysis of different compositions GelMA-HAMA hydrogel (5% GelMA / 2% HAMA (5G/2H); 3% GelMA / 1% HAMA (3G/1H); 2.5% GelMA / 2.5% HAMA (2.5G/2.5H); and 2% GelMA / 1% HAMA (2G/1H)): **B**. Storage modulus (G’) and **C**. Loss modulus (G”). Temperature weep analysis of 3GH1 hydrogel crosslinked for 5 s, 15 s, and 20 s: **D**. Storage modulus (G’) and **E**. Loss modulus (G’). Frequency sweep analysis of 3GH1 hydrogel crosslinked for 5 s, 15 s, and 20 s: **F**. Storage modulus (G’) and **G**. Loss modulus (G”). Amplitude sweep analysis of 3GH1 hydrogel crosslinked for 5 s, 15 s, and 20 s: **H**. Storage modulus (G’) and **I**. Loss modulus (G”). **J**. Swelling study of 3GH1 hydrogel up to 48 hours. **K**. Degradation study of 3GH1 hydrogel up to 7 weeks. **L**. SEM analysis of 3GH1 hydrogel crosslinked for 5 s, 15 s, and 20 s (Scale bars: 50 μm). **M**. Porosity analysis of 3GH1 hydrogel crosslinked for 5 s, 15 s, and 20 s.

### 2.2. Characterization of GelMA-HAMA-based GBM bioconstructs

We fabricated GelMA-HAMA-based 3D GBM bioconstructs using three human GBM cell lines: LN229, PDM140, and U87. We generated GBM spheroids using agarose microwell arrays, and we monitored the formation of spheroids daily for 3 days and characterized their viability and morphology (area, aspect ratio, and circularity). Before encapsulation in the 3G/1H hydrogel, LN229 spheroids exhibited an average viability of 95.6%, area of 24488 μm2, aspect ratio of 1.01, and circularity of 0.47; PDM140 spheroids exhibited an average viability of 96.5%, area of 30007 μm2, aspect ratio of 1.27, and circularity of 0.49; and U87 spheroids exhibited an average viability of 98.3%, area of 32302 μm2, aspect ratio of 1.24, and circularity of 0.52 (**Figure S3**). Next, we encapsulated the GBM spheroids in the 3G/1H hydrogel to fabricate 3D GBM bioconstructs recapitulating the mechanical properties of brain tissues. We characterized the viability and morphology of the different GBM bioconstructs on days 1, 7, and 14 of culture. On day 14, the viability of LN229, PDM140, and U87 cells remained above 95.9%, 94.8%, and 94.8%, respectively (**Figures 2A and 2B**). The morphological analysis of the GBM spheroids encapsulated in the 3G/1H hydrogel revealed that the area of LN229, PDM140, and U87 spheroids significantly increased from 35600 μm2 to 72000 μm2, 15500 μm2 to 68700 μm2, and 14000 μm2 to 53200 μm2, respectively, between day 1 and 14 of culture (**Figure 2C**). Similarly, the aspect ratio of LN229, PDM140, and U87 spheroids increased from 1.08 to 1.28, 1.1 to 1.27, and 1.05 to 1.35, respectively, between days 1 and 14 of culture (**Figure 2D**). Conversely, the circularity of LN229, PDM140, and U87 spheroids significantly decreased from 0.35 to 0.05, 0.31 to 0.04, and 0.37 to 0.04, respectively, between days 1 and 14 of culture (**Figure 2E**). The observed viability and morphological changes demonstrated that the 3D GBM bioconstructs, using human LN229, PDM140, and U87 GBM cells, were viable and proliferative for 14 days in our 3G/1H hydrogel mimicking the mechanical properties of brain tissues.

**Figure 2:**
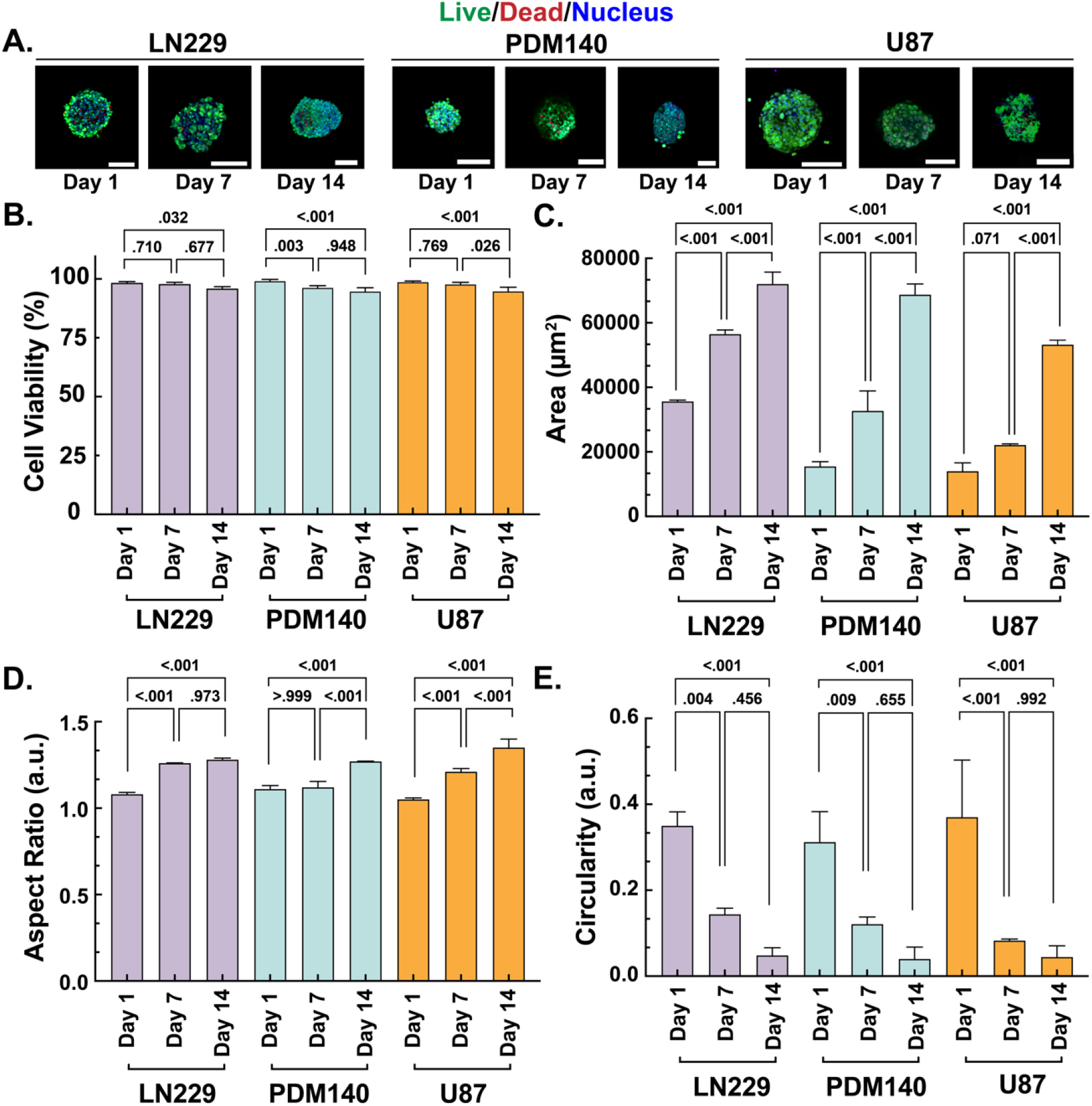
Characterization of large GelMA-HAMA-based 3D GBM models using LN229, PDM140, and U87 cells after 1, 7, and 14 days of culture. **A**. Live (green) and dead (red) staining of LN229, PDM140, and U87 spheroid encapsulated in the 3G/1H hydrogel (scale bar = 150 μm). **B**. Viability of LN229, PDM140, and U87 spheroids. **C**. Area of LN229, PDM140, and U87 spheroids. **D**. Aspect ratio of LN229, PDM140, and U87 spheroids. **E**. Circularity of LN229, PDM140, and U87 spheroids.

### 2.3. TMZ sensitivity assessment for LN229, PDM140, and U87 GBM bioconstructs

We used our 3D GBM bioconstructs to assess the sensitivity of LN229, PDM140, and U87 spheroids to TMZ. We observed that the viability of all spheroids tested exhibited a dose-dependent response to TMZ (**Figure 3A**). The viability of LN229 spheroids was significantly lower than the control at a TMZ concentration of 100 μM (p<0.05), and at a concentration of 1000 μM, the viability of LN229 spheroids was 8.52% (**Figure 3B**). The viability of PDM140 spheroids was significantly lower than the control at a TMZ concentration of 50 μM (p<0.001), and at a concentration of 100 μM, the viability of LPDM140 spheroids was 30.4% (**Figure 3C**). Beyond 100 μM, the viability of PDM140 spheroids did not decrease further. The viability of U87 spheroids was significantly lower than the control at a TMZ concentration of 250 μM (p<0.01), and at a concentration of 1000 μM, the viability of U87 spheroids was 26.0% (**Figure 3D**). The viability of GBM spheroids was confirmed using confocal analysis, where the viability of LN229 was 4.25% at a concentration of 1000 μM of TMZ, the viability of PDM140 was 29.8% at a concentration of 100 μM of TMZ, and the viability of U87 was 29.2% at a concentration of 1000 μM of TMZ (**Figure S4A-C**). The morphological analysis of the GBM spheroids revealed that the area and circularity of LN229, PDM140, and U87 spheroids significantly decreased upon TMZ treatment (p<0.001) (**Figures 3E and 3F**). In addition, the aspect ratio of LN229, PDM140, and U87 spheroids significantly changed upon TMZ treatment (p<0.001) (**Figure 3G**). Lastly, the dose-response curves revealed that PDM140 spheroids were most sensitive to TMZ (IC_50_= 73 μM), followed by LN229 (IC_50_= 278 μM) and U87 (IC_50_= 446 μM) (**Figure 3H**). The Area Under the Curve (AUC) further confirmed the difference in TMZ sensitivity between the different GBM spheroids (p<0.001), with PDM140 being the most sensitive, followed by LN229 and U87 being the most resistant to TMZ (**Figure 3H**).

**Figure 3:**
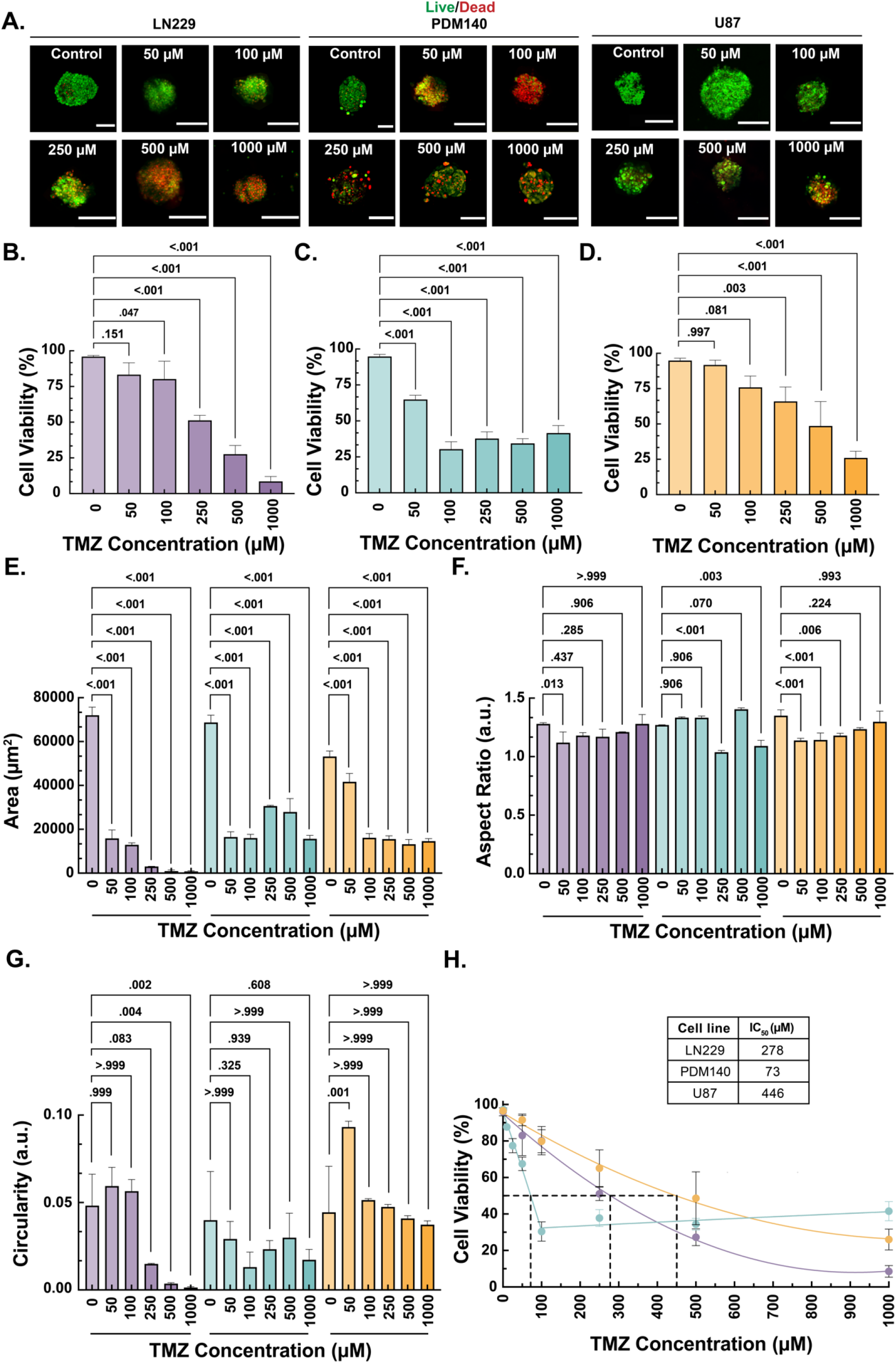
TMZ sensitivity assessment using LN229, PDM140, and U87 GBM bioconstructs. **A**. Live (green) and dead (red) staining of LN229, PDM140, and U87 spheroid with increasing concentrations of TMZ (scale bar = 150 μm). **B**. Viability of LN229 spheroids in response to different concentrations of TMZ. **C**. Viability of PDM140 spheroids in response to different concentrations of TMZ. **D**. Viability of U87 spheroids in response to different concentrations of TMZ. **E**. Area of LN229, PDM140, and U87 spheroids in response to different concentrations of TMZ. **F**. Aspect ratio of LN229, PDM140, and U87 spheroids in response to different concentrations of TMZ. **G**. Circularity of LN229, PDM140, and U87 spheroids in response to different concentrations of TMZ. **H**. TMZ Dose-response curves with IC_50_ values and Area Under the Curve analysis for LN229, PDM140, and U87.

### 2.4. Characterization and TMZ sensitivity assessment for human pericyte bioconstructs

We fabricated and characterized GelMA-HAMA-based 3D human pericyte bioconstructs. We encapsulated the pericytes in the 3G/1H hydrogel and assessed their viability after days 1, 7, and 14 of culture (**Figure 4A**). The viability of the pericytes remained above 95% after 14 days of culture (**Figure 4B**). Next, we treated the pericyte bioconstructs with different concentrations of TMZ (0-1000 μM) on day 7 of culture. On day 14, we observed that the viability of the pericytes exhibited a dose-dependent response to TMZ (**Figure 4C**). The viability of the pericytes was significantly lower than the control at a TMZ concentration of 50 μM (p<0.05), and at a concentration of 1000 μM, the viability of the pericytes was 44.6% (**Figure 4D**). The viability of the pericytes was confirmed using confocal analysis, where, at a concentration of 1000 μM of TMZ, their viability was 42.8% (**Figure S4D**). The dose-response curve showed that the IC_50_ value of TMZ on pericytes was 790 μM (**Figure 4E**). Lastly, the AUC of TMZ treatment on the primary pericytes (AUC=61405) was higher than the AUC for LN229 (AUC=34684), PDM140 (AUC=6263), and U87 (AUC=51403).

**Figure 4:**
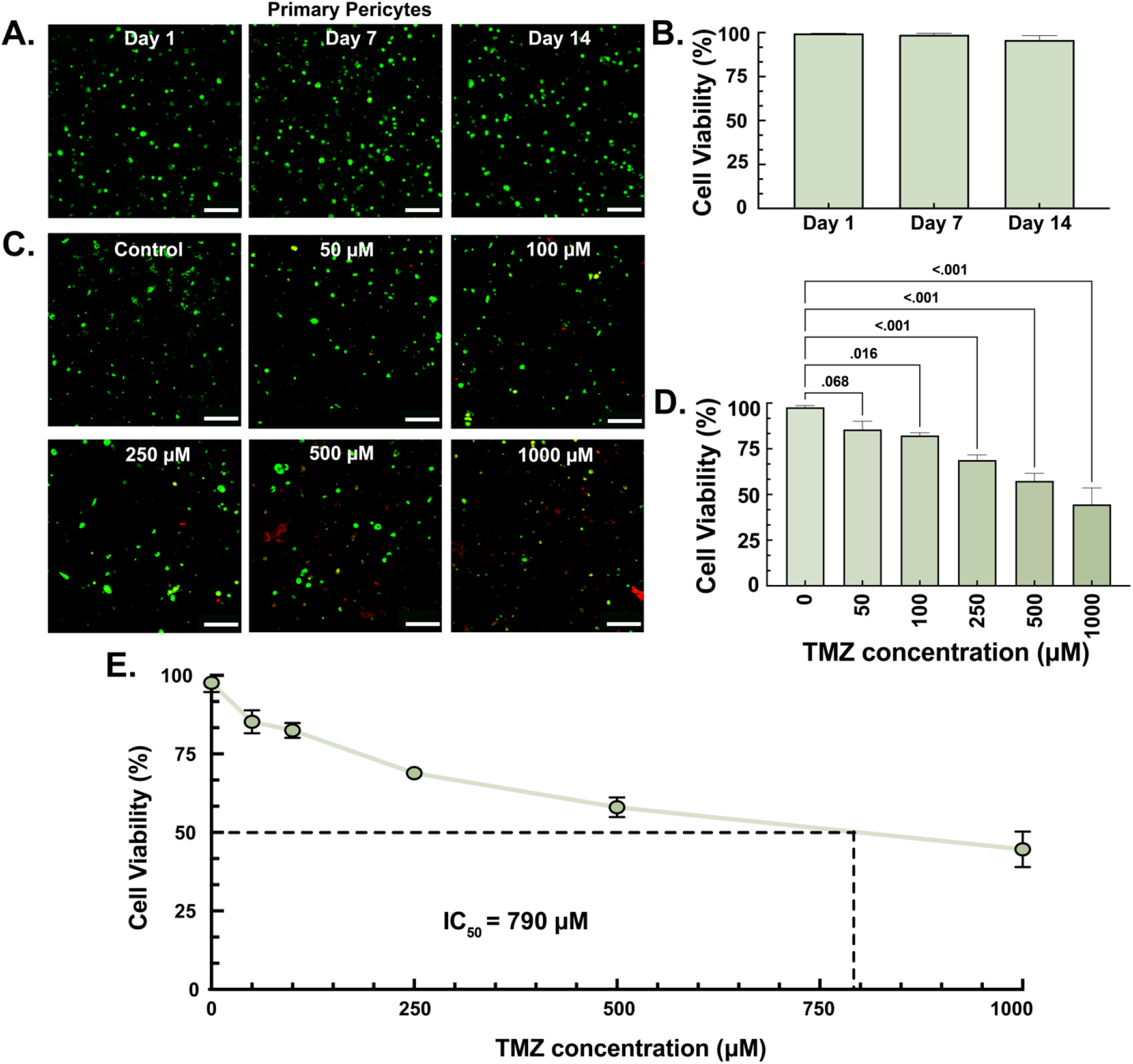
Characterization of a large GelMA-HAMA-based 3D primary pericytes model and TMZ sensitivity assessment. **A**. Live (green) and dead (dead) staining of primary pericytes encapsulated in the 3G/1H hydrogel after 1, 7, and 14 days of culture (scale bar = 150 μm). **B**. Viability of primary pericytes after 1, 7, and 14 days of culture. **C**. Live (green) and dead (staining) of primary pericytes with increasing concentrations of TMZ (scale bar = 150 μm). **D**. Viability of primary pericytes in response to different concentrations of TMZ. **E**. TMZ dose-response curves with IC_50_ values and AUC for primary pericytes models.

### 2.5. Gene expression analysis in GBM or pericyte bioconstructs upon TMZ treatment

The expression of GADD45a, PTEN, MSH6, and CCL5 in human GBM spheroids (LN229, PDM140, and U87) or pericytes bioconstructs was evaluated using qRT-PCR on days 7 and 14 of culture. For LN229 bioconstructs: the relative mRNA expression of GADD45a,

PTEN, and MSH6 was 3.68, 3.94, and 0.06 on day 7 and significantly increased to 14.0, 89.0, and 18.1 on day 14 (p<0.001), respectively, and the relative mRNA expression of CCL5 was 1.42 on day 7 and did not significantly increase on day 14 (3.30 relative mRNA expression) (**Figure 5A**). When LN229 bioconstructs were treated with 1000 μM of TMZ, the relative mRNA expressions of GADD45a and PTEN on day 14 were significantly downregulated compared to untreated bioconstructs (14.0 vs. 1.31 (p<0.001) and 89.0 vs. 22.5 (p<0.001), respectively), and no significant expression changes were observed for MSH6 and CCL5 (**Figure 5A**). For PDM140 bioconstructs: the relative mRNA expressions of GADD45a, MSH6, and CCL5 were 1.54, 0.69, and 1.53 on day 7, respectively, and did not significantly increase on day 14 (2.30, 1.48, and 0.43 relative mRNA expression, respectively), whereas the relative mRNA expression of PTEN was 8.24 on day 7, and significantly increased to 98.5 on day 14 (p<0.001 (**Figure 5B**). When PDM140 bioconstructs were treated with 100 μM of TMZ, the expression of PTEN was significantly downregulated compared to untreated bioconstructs (98.5 vs. 33.0, p<0.001) whereas the expression of MSH6 was significantly upregulated (2.73 vs. 8.74, p<0.001), and no significant expression changes were observed for GADD45a and CCL5 (**Figure 5B**). For U87 bioconstructs, the relative mRNA expressions of GADD45a, PTEN, MSH6, and CCL5 were 0.46, 1.07, 1.40, and 1.12 on day 7, respectively, and did not significantly increase on day 14 (0.99, 1.63, 1.48, and 1.04 relative mRNA expression, respectively) (**Figure 5C**). When U87 bioconstructs were treated with 1000 μM of TMZ, the expressions of PTEN and MSH6 were significantly upregulated compared to untreated bioconstructs (1.63 vs. 275 (p<0.001) and 1.48 vs. 29.6 (p< 0.001), respectively), and no significant expression changes were observed for GADD45a and CCL5 (**Figure 5C**). For pericytes bioconstructs, the relative mRNA expressions of GADD45a, PTEN, MSH6, and CCL5 were 1.01, 2.9, 0.06, and 0.06 on day 7, respectively, and did not significantly increase on day 14 (1.09, 2.78, 0.45, and 0.43relative mRNA expression, respectively) (**Figure 5D**). When pericyte bioconstructs were treated with 1000 μM of TMZ, the expressions of GADD45a, PTEN, MSH6, and CCL5 expressions were significantly upregulated compared to untreated bioconstructs (1.9 vs. 10.1 (p<0.001), 2.78 vs. 8.75 (p<0.001), 0.45 vs. 105 (p<0.001), and 0.42 vs. 163 (p<0.001), respectively) (**Figure 5D**).

**Figure 5:**
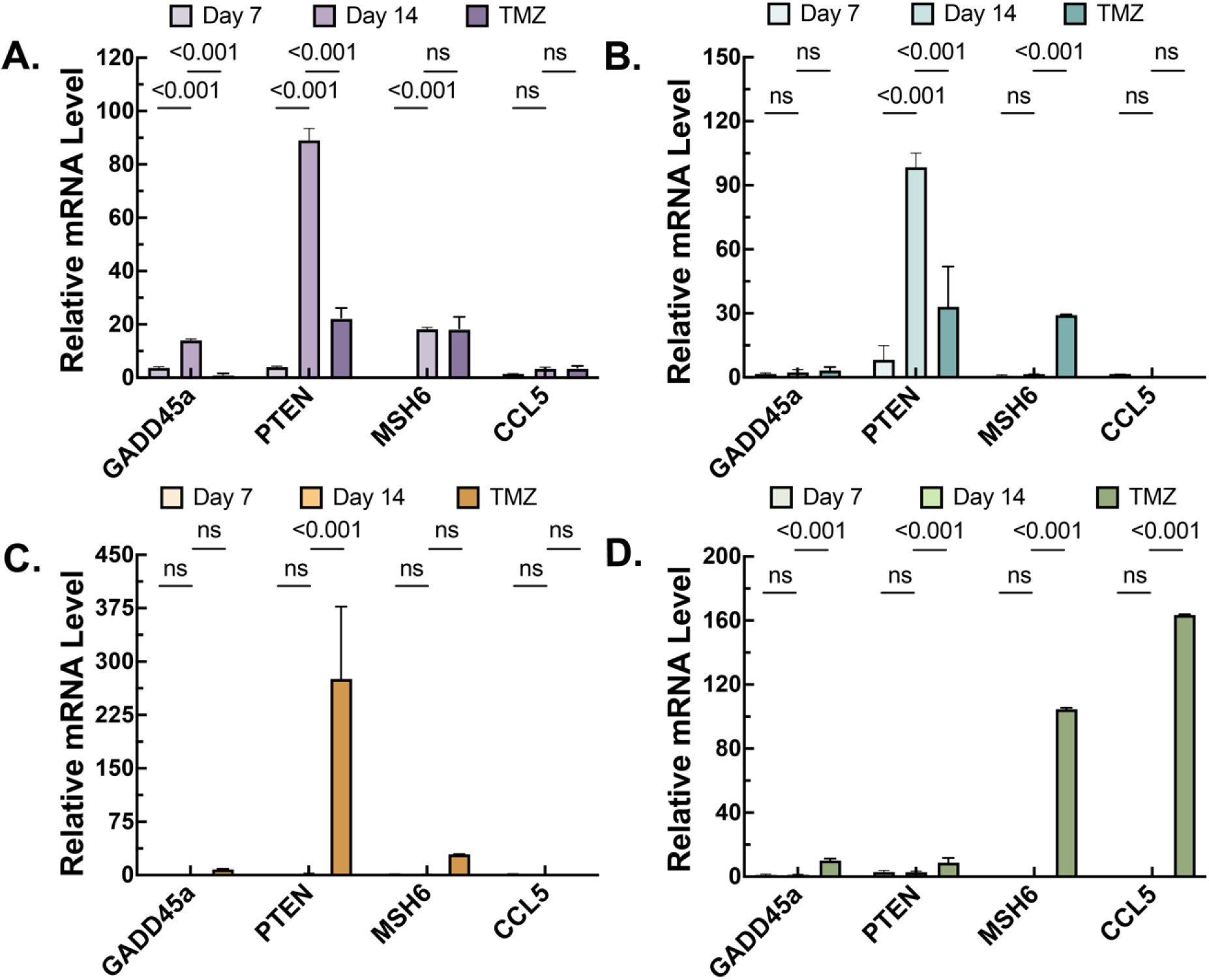
qPCR analysis on LN229, PDM140, U87, and Pericytes. Change in the relative mRNA expression for GADD45a, PTEN, MSH6, and CCL5 in **A**. LN229, **B**. PDM140, **C**. U87, and **D**. pericytes.

### 3.6. Characterization and TMZ sensitivity assessment for multicellular GBM/pericytes bioconstructs

We fabricated and characterized GelMA-HAMA-based 3D multicellular GBM/pericytes bioconstructs to evaluate GBM TMZ resistance induced by pericytes. Within the GBM/pericyte bioconstructs, we observed that the pericytes were located on the surface of the GBM spheroids and their surroundings (**Figure 6A-D**). The average viability of the LN229/pericytes, PDM140/pericytes, and U87/pericytes bioconstructs was assessed on days 1, 7, and 14 using confocal image analysis. After 14 days of co-culture without TMZ treatment, the viability of the LN229, PDM140, and U87 spheroids was 91.8%, 89.3%, and 84.8%, respectively. The viability of the corresponding pericytes was 89.5%, 85.2%, and 81.0%. In contrast, for TMZ-treated GBM/pericytes bioconstructs, the viability of the LN229, PDM140, and U87 spheroids was 41.0%, 53.1%, and 48.1%, respectively, and the viability of the corresponding pericytes was 66.6%, 64.9%, and 55.8% (**Figure 6E**). When comparing the viability of TMZ-treated LN229, PDM140, and U87 spheroids with and without pericytes, we observed that the viability of GBM spheroids was higher in the presence of pericytes (**Table 1**). LN229 spheroids treated with 1000 μM of TMZ displayed significantly higher viability in the presence of pericytes than in the absence of pericytes (41.0% vs. 8.52%, p<0.001). PDM140 spheroids treated with 100 μM of TMZ displayed significantly higher viability in the presence of pericytes than in their absence (53.1% vs. 30.4%, p<0.001). U87 spheroids treated with 1000 μM of TMZ displayed significantly higher viability in the presence of pericytes than in their absence (48.1% vs. 26.0%, p<0.001).

**Table 1:**
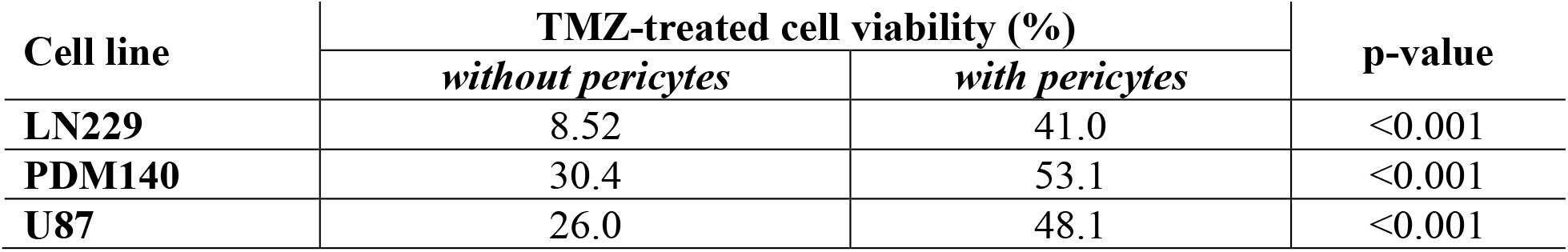
Cell viability, IC_50_ values, and p-values in mono-and co-culture systems.

**Figure 6:**
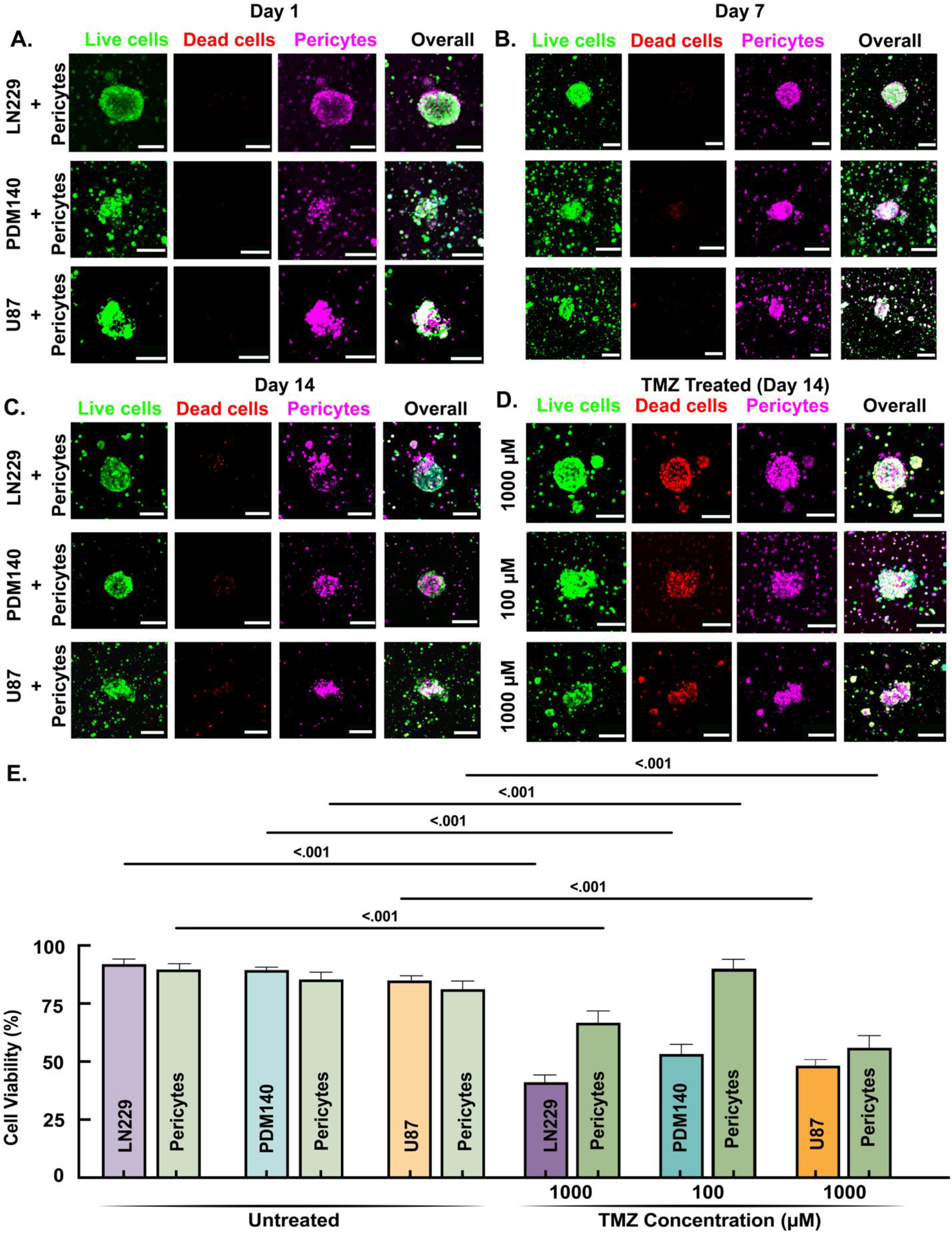
TMZ sensitivity assessment using GBM/pericytes bioconstructs. Live (green), dead (red), and pericytes (magenta) staining of LN229, PDM140, and U87/pericytes bioconstructs on **A**. day 1, **B**. day 7, **C**. day 14 (untreated), and **D**. day 14 (TMZ-treated) (scale bar = 150 μm). **E**. Day 14 cell viability of GBM cells (LN229, PDM140, and U87) and pericytes in GBM/pericyte bioconstructs with and without TMZ treatment.

## 3. Discussion

We developed a novel multicellular 3D GBM MPS mimicking the physicochemical properties of brain tissues and pericyte-mediated TMZ resistance that can be used to screen for standard-of-care chemotherapy (i.e., TMZ). This MPS comprised GBM spheroids and pericytes encapsulated in a GelMA-HAMA-based hydrogel that mimics the effect of the interaction between the GBM cells and the TME on TMZ sensitivity. Using this 3D GBM MPS, we demonstrated: (1) the differential sensitivity to TMZ in several GBM cell lines (with PDM140 being the most sensitive, followed by LN229 and U87); (2) the off-target toxicity of TMZ towards pericytes; and (3) pericyte-induced TMZ resistance in across several GBM cell lines. These results confirm the important role of pericytes in GBM resistance to TMZ, possibly mediated through the CCL5-CCR5 paracrine axis.^[18]^ Indeed, upon TMZ treatment, we observed that pericytes drastically overexpressed CCL5 (380-fold increase), and, in the presence of pericytes, GBM cells demonstrated a 22-33% increase in TMZ resistance compared to GBM cells culture without pericytes. These findings demonstrate the impact of the complex interactions between GBM cells, pericytes, and the TME on chemoresistance, thus emphasizing the importance of including different components of the GBM TME to assess drug sensitivity profiles.

Our novel 3D GBM MPS that closely mimics the physicochemical properties of brain tissue could significantly advances our understanding of GBM and its resistance to TMZ. Traditional two-dimensional (2D) cell cultures have been widely used in GBM research, yet they fail to accurately replicate the complex GBM TME, leading to discrepancies in drug sensitivity and resistance observed in clinical settings.^[30]^ Previous studies have utilized various 3D culture models, including spheroids, which provide a more physiologically relevant environment for tumor cells.^[21]^ However, these models often lack crucial stromal components such as pericytes, which play an essential role in the GBM TME.^[31]^ Incorporating pericytes into the 3D GBM MPS used in this study offers a substantial improvement, as it allows for examining pericyte-mediated mechanisms of drug resistance, an underexplored area.^[15]^ This study ‘s findings demonstrate the importance of including diverse cellular components of the TME in GBM models to accurately represent the *in vivo* conditions, thereby enhancing the reliability and relevance of drug testing.^[32]^ Furthermore, the developed 3D GBM MPS provides a robust platform for investigating the interactions between GBM cells and pericytes, revealing critical insights into the mechanisms underlying chemoresistance and highlighting the potential for novel therapeutic interventions targeting the CCL5-CCR5 paracrine axis. By comparing the outcomes of this study with those of previous research, it becomes evident that the inclusion of pericytes in the 3D model offers a more comprehensive understanding of the TME and its impact on TMZ resistance in GBM.

Hydrogels are an integral part of 3D GBM MPS as they provide a supportive matrix that mimics the ECM of brain tissues and provides specific properties that enhance various aspects of tumor modeling.^[33]^ Several types of hydrogels have been used, each selected for particular functions based on their unique properties. Matrigel, the most popular hydrogel matrix used for GBM modeling *in vitro*, is rich in ECM proteins (laminin, collagen IV, and heparan sulfate proteoglycans) and provides a highly biomimetic environment that promotes cell adhesion, growth, and differentiation.^[34]^ Other types of hydrogels that have been used to create 3D GBM models include collagen, alginate, polyethylene glycol (PEG), fibrin, HA, and GelMA, which are generally selected for their biocompatibility, functionality, tunable mechanical properties, and ability to support cell attachment and migration, all of which are essential properties that are beneficial to study tumor behaviors (i.e., cell proliferation, migration, differentiation, and invasion) and drug responses. Nonetheless, the mechanical and rheological properties of these hydrogels are mismatched with physiological brain tissue properties despite their essential role during neurodevelopment and homeostasis.^[26]^

The selection of a GelMA-HAMA-based composite hydrogel allowed us to generate a scaffold with rheological properties within the range of brain tissues (300<G’<1200Pa and 20<G”<120Pa),^[29]^ which is not achievable with GelMA alone.^[35]^ This hydrogel composite combines the robust structural support of GelMA with the bioactive and hydrophilic nature of HA, resulting in a hydrogel with improved viscoelastic properties, tunable mechanical strength, and enhanced biocompatibility. As described previously, the physiochemical properties of our GelMA-HAMA hydrogel remained relatively unchanged during long-term (14 days) cell culture conditions while exhibiting slow degradation rates typical of GelMA despite the faster degradability of HAMA and being sufficiently porous to allow the rapid diffusion of small molecules, nutrients, and oxygen throughout large bioconstructs (98 mm^3^).^[36]^ Also, the synergistic effects of GelMA and HAMA ensure consistent gelation and structural integrity under physiological conditions. Overall, our 3D GBM MPS is a promising platform for advanced GBM research and therapeutic testing, which includes many important factors for replicating the dynamic mechanical environment of brain tissues, thereby providing a more accurate model for studying GBM mechanics and cellular interactions within the ECM.^[37]^

Numerous *in vitro* studies have assessed TMZ sensitivity in different GBM cell lines, primarily using 2D culture systems.^[38]^ Recent 3D GBM models, either as spheroids of GBM cell encapsulated in a hydrogel matrix, demonstrated more physiological relevance due to enhanced cell-cell and cell-matrix interactions activating cellular pathways showing drug resistance and outer spheroids layer acting as a physical barrier to TMZ hindering diffusion to the core.^[39]^ Our 3D GBM MPS combines the advantages of spheroids and the presence of brain-like ECM, mimicking the diffusion of TMZ after crossing the BBB into brain/GBM tissues. Indeed, the IC_50_ values of TMZ sensitivity for LN229 and U87 in 2D cultures were approximately 110 μM and 220 μM, respectively;^[6c, 38]^ however, the IC_50_ values for LN229 and U87 spheroids encapsulated in our GelMA-HAMA hydrogel were higher, suggesting increased resistance to TMZ resulting from penetration and diffusion of the drug.

The differential sensitivity of GBM cell lines, with or without pericytes, to TMZ demonstrates the complexity of cell-specific chemoresistance mechanisms. Notably, PDM140 cells were the most sensitive to TMZ, followed by LN229 and U87. The role of pericytes in enhancing GBM spheroid viability under TMZ treatment confirms their active involvement in promoting TMZ resistance.^[18]^ Indeed, pericytes promote the growth of cancer cells and drug resistance through paracrine signaling by secreting various cytokines and growth factors.^[17]^ A key mechanism implicated in pericyte-induced chemoresistance is the CCL5-CCR5 paracrine axis. Pericytes secrete the chemokine CCL5, which binds to the CCR5 receptor on GBM cells, activating signaling pathways that activate the DNA damage response upon TMZ treatment and promote cell survival, proliferation, and resistance to apoptosis. This paracrine signaling can enhance the invasive capabilities of GBM cells and their resistance to TMZ by activating downstream pathways associated with tumor progression, such as PI3K/AKT and MAPK.^[40]^ Moreover, the CCL5-CCR5 axis facilitates the recruitment of additional stromal cells to the GBM TME, further reinforcing the pro-survival conditions that contribute to chemoresistance.^[41]^

Upon TMZ treatment, we observed a significant upregulation of CCL5 (380-fold increase) in pericytes only, confirming previous findings demonstrating the contribution of pericyte-secreted CCL5 to the resistance of GBM cells to TMZ.^[18]^ Our results indicate that the spheroids do not disrupt the CCL5-CCR5 paracrine axis in the presence of a hydrogel matrix. Furthermore, the modulation of the expression of TMZ resistance-associated genes (i.e., GADD45a, PTEN, and MSH6) also contributes to the observed differential sensitivity of GBM cell lines, with or without pericytes, to TMZ. Indeed, we observed differential expression of GADD45a, PTEN, and MSH6 in different GBM cells upon TMZ treatment. The most TMZ-resistant cell line, U87, had a defective expression of GADD45a with a significant increase in PTEN expression; LN229 significantly downregulated GADD45a and PTEN expression; and the most sensitive cell line, PDM140, significantly downregulated PTEN and upregulated MHS6. These observations align with the current understanding of gene expression modulation and TMZ resistance, where the loss or decreased expression of GADD45a, PTEN, and MSH6 is associated with TMZ resistance^.[42]^ Our study confirms that GBM cells and components of the TME, including pericytes, can significantly modulate the response to TMZ. Importantly, modeling the CCL5-CCR5 paracrine axis using our 3D GBM MPS represents an ideal in vitro platform to test novel therapeutic interventions aimed at disrupting this chemoresistance mechanism and improving the efficacy of TMZ treatment in GBM patients.

Despite the advancements presented in our study, several limitations must be acknowledged. First, our 3D GBM model does not encompass the full complexity of the GBM TME, lacking critical ECM components (i.e., fibronectin and laminin) and other cell types (i.e., astrocytes, microglia, and immune cells) that contribute to TMZ resistance.^[43]^ Second, our 3D GBM model of TMZ resistance is limited to the CCL5-CCR5 axis and intrinsic GBM mutations, but other resistance mechanisms crucial for immune evasion (e.g., PD-L1 and CD47 upregulation)^[44]^ represent important aspects of the TMZ resistance landscape that are absent in our model. Third, the treatment regimen applied in this study spans only seven days, which does not accurately reflect the extended treatment schedules used in clinical settings. Clinical regimens for TMZ often involve multiple treatment cycles over several weeks to months, which could influence the development and observation of resistance mechanisms over time.^[1b, 1c]^ Fourth, using commercially available cell lines, such as PDM140, LN229, and U87, rather than patient-derived cells limits the clinical relevance of the findings. Commercial cell lines may not fully capture the genetic and phenotypic heterogeneity of patient tumors, thus impacting the translational applicability of the results.^[45]^ Lastly, our model lacks critical components of the brain environment, such as a functional BBB. The absence of the BBB in the model is particularly limiting, as it plays a significant role in drug delivery and efficacy. Drugs that appear effective in this model may not necessarily cross the BBB *in vivo*, leading to potentially misleading results regarding their clinical relevance.^[46]^ Furthermore, the lack of vasculature precludes the assessment of angiogenesis and the interaction between tumor cells and blood vessels, which are essential for understanding tumor growth and metastasis.^[47]^

## 4. Conclusions

Our study demonstrates the development and application of a novel multicellular 3D GBM MPS model designed to replicate the physicochemical properties of brain tissues and the pericyte-mediated resistance to TMZ. This advanced hydrogel-based platform demonstrated the critical role of the GBM TME in modulating chemotherapy sensitivity, particularly emphasizing the influence of pericytes in promoting TMZ resistance. The differential drug sensitivities observed among various GBM cell lines illustrate the necessity for patient-specific models to facilitate personalized treatment strategies and improve GBM patient outcomes. A significant benefit of this study is the modeling of the pericyte-induced CCL5-CCR5 axis, which constitutes an ideal platform enabling a better understanding of the paracrine signaling mechanisms contributing to TMZ resistance in GBM. Therefore, including this paracrine axis allows for a more accurate simulation of the TME and potentially leads to the identification of novel therapeutic targets within the CCL5-CCR5 pathway toward more effective treatments disrupting these resistance mechanisms. While our 3D GBM MPS model provides a valuable platform for studying GBM and TMZ resistance, its limitations confirm the need for more comprehensive models that better replicate the intricate TME, include a broader range of resistance mechanisms, adopt clinically relevant treatment regimens, utilize patient-derived cells, and incorporate critical physiological barriers such as the BBB. Overall, the proposed 3D GBM MPS model can transform drug screening and personalized treatment for GBM by offering ethical and cost-effective alternatives to animal testing and more effective drug screening and discovery efforts, ultimately improving GBM patient outcomes.

## 5. Experimental Section/Methods

### GelMA Synthesis

The gelatin was subjected to methacryloyl modification following our previously published procedure.^[36a]^ Type A porcine gelatin (300 bloom, Sigma G1890) was dissolved in Dulbecco’s phosphate-buffered saline (DPBS; GIBCO) at 50°C to produce 10% (w/v) gelatin. Methacrylic anhydride (MAA) was gradually incorporated into the gelatin solution using a syringe pump dispensing 0.5 mL/min to achieve a 2X molar excess. After adding MAA, the reaction was maintained at 50°C with continuous stirring for an hour. The reaction was then halted by incorporating an equivalent volume of DPBS and adjusting the pH to 7.4. Upon completion of the methacrylation process, the samples were centrifugated, and the supernatants were dialyzed against distilled water. Following the dialysis, the aqueous reaction solution was freeze-dried to produce GelMA.

### HAMA Synthesis

The methacryloyl modification of HA was conducted based on previously published procedures.^[48]^ To prepare a 1% (w/v) HA solution, a dry powder form of sodium hyaluronate (750 kDa - 1 MDa, LifeCore) was dissolved in distilled water. In order to decrease viscosity and enhance the miscibility of MAA, dimethylformamide (DMF) was added to the solution (DMF:water = 1:1), and the solution was stirred on the ice bath and cooled down to 4°C. Meanwhile, the 3x molar excess MAA was gradually added to the HA solution drop by drop. As a result, the hydroxyl residues of the N-Acetylglucosamine (GlcNAc) repeating unit of the polymer backbone were methacrylated. By continuously monitoring and adding 1 M NaOH, the pH of the reaction was maintained at 9 throughout the process. The reaction lasted 6h to stabilize the pH, and the solution was stored on ice overnight. The components were moved to a dialysis tubing the next day and were dialyzed in distilled water for a week. Following dialysis, the samples underwent centrifugation, and the resultant supernatant was freeze-dried to produce HAMA.

### NMR Characterization

Proton NMR was performed to confirm that the methacrylation of HA and gelatin was successful. Proton NMR was carried out with 16 scans at a frequency of 400 MHz. Deuterium oxide (D_2_O, 20 mg/mL) was used to dissolve the HAMA and GelMA hydrogel.

### Preparation of GelMA-HAMA Network Hydrogels

We synthesized four different composite hydrogels dissolved in cell culture medium with 1% (w/v) Irgacure (Irgacure, 2-Hydroxy-4′-(2-hydroxyethoxy)-2-methylpropiophenone): 5% GelMA / 2% HAMA (5G/2H); 3% GelMA / 1% HAMA (3G/1H); 2.5% GelMA / 2.5% HAMA (2.5G/2.5H); and 2% GelMA / 1% HAMA (2G/1H). First, we dissolved the photoinitiator in cell culture media at 50°C with constant stirring. Next, HAMA was incorporated into the solution and dissolved at 4°C overnight. Last, GelMA was incorporated and dissolved at 50°C with continuous stirring to generate the GelMA-HAMA prepolymer solution. To generate a cylindrical hydrogel construct (diameter: 5 mm; and height: 5 mm), we cast 98 μL of prepolymer solution in a cylindrical shell PMMA mold (inner diameter: 5 mm; outer diameter: 12 mm; and height: 6 mm), and the prepolymer solution was crosslinked with UV for 5, 15, 20, 30, or 60 seconds using a 365 nm point light source (Omnicure 2000) with a power intensity of 500 mW/cm^2^. The resulting hydrogel constructs were gently removed from the PMMA mold and used in subsequent physicochemical characterization experiments.

### Hydrogel Characterization

#### (i) Fourier Transform Infrared - Attenuated Total Reflection (FTIR-ATR)

The FTIR spectra of the GelMA, HAMA, and GelMA-HAMA were obtained using FTIR (Bruker, Alpha II, Germany). The samples were scanned over a wavenumber ranging from 4000 to 400 cm^−1^, employing a resolution of 4 cm^−1^.

#### (ii) Rheology

The rheological characteristics (storage modulus (G’) and loss modulus (G”)) of HAMA, GelMA, and HAMA-GelMA hydrogels were measured in terms of temperature sweep, frequency sweep, and amplitude sweep using a Rheometer (Anton Paar, MCR 302) with a 25 mm sandblasted parallel plate geometry. Frequency sweeps were performed in the range of 0.1 to 10 Hz at a constant strain of 1% and a temperature of 37°C. Temperature sweeps were performed at a frequency of 1 Hz, a constant strain of 1%, and a temperature ranging from 24 to 45°C. Amplitude sweeps were performed in the range of 8.9×10^−5^ to 1 % at a frequency of 1 Hz and a temperature of 37°C. The Anton Paar Rheocompass software was used to record the storage and loss moduli data.

#### (iii) Swelling

For swelling experiments, GelMA-HAMA hydrogels were immersed in 1X PBS. At different time points (0 min, 5 min, 10 min, 30 min, 60 min, 24 hr, and 48 hr), the excess PBS on the surface of the hydrogels was removed using dry tissue paper. The hydrogel weight was measured, where W_i_ is the initial weight of the hydrogels, and W_n_ is their weight after water uptake at t=n.^[49]^ The swelling rates were calculated with equation (1):

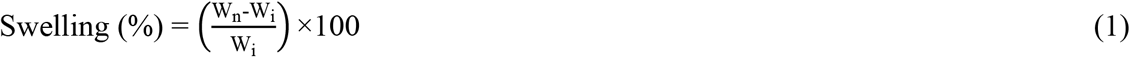

#### (iv) Degradation

For the degradation study, all GelMA-HAMA hydrogel samples were freeze-dried for 1 day before the initial weight (W_0_) was measured. Next, the hydrogels were placed inside an incubator at 37ºC and soaked in 1X PBS. Every week until week 7, the hydrogel samples were removed from the 1X PBS and vacuum-dried at 50°C. The final dry weight (W_t_) of the samples was recorded before being soaked again in 1X PBS. The degradation rates were determined utilizing the following equation (2):^[50]^

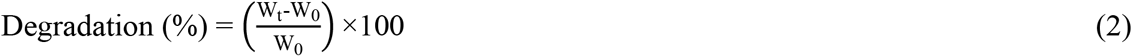

#### (v) Porosity Analysis

The porosity analysis was performed using SEM images of the GelMA-HAMA hydrogel samples. The samples were lyophilized, fixed to the SEM stubs, and vacuum coated with gold for 30 s. The samples were visualized using a Zeiss Supra V 40VP SEM under high vacuum conditions. The resulting images were analyzed using the ImageJ software (http://rsbweb.nih.gov/ij/) and the Fiji plug-in to determine pore sizes. The SEM images of the hydrogel were changed to 8-bit grayscale before adjusting the threshold values to determine the boundaries of the pores in the hydrogel. The total area of the pores (AP) and the total area of the SEM image (TI) were measured using the image volume method.^[51]^ Finally, the porosity was calculated with equation (3):

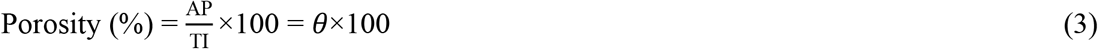

### Simulation of TMZ molecule diffusion in GelMA-HAMA hydrogels

A 2D computational model was developed in COMSOL Multiphysics using the finite element-based method to simulate the diffusion of TMZ molecules in porous GelMA-HAMA hydrogels immersed in an aqueous solution. An axis-symmetric model was developed using “creeping flow” and “transport of diluted species in porous media” equations with the specific dimension of our hydrogel constructs (radius: 2.5 mm; and height: 5 mm) and surrounding media (height and width: 12 mm) (Figure S1A). The hydrogel and the media were set as isothermal, incompressible, and Newtonian. The governing equations for this model are as follows:^[52]^

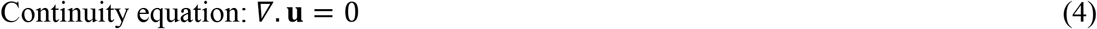

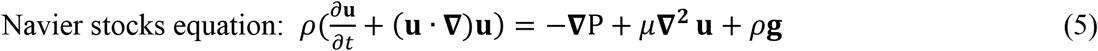

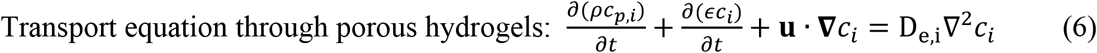

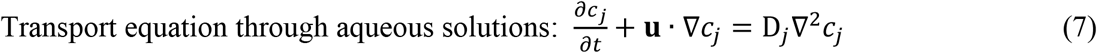

The symbols **u**, *ρ, μ*, **g**, P, *ϵ, c*_*i*_, *c*_*j*_, *c*_*p,i*_, D_e,i_, and D_*j*_ refer to velocity, the density of the water, dynamic viscosity, acceleration due to gravity, pressure, the porosity of hydrogel, concentration of TMZ in the media inside the pores, concentration of TMZ in the media outside the hydrogel, the concentration of TMZ in the solid hydrogel, diffusion co-efficient of TMZ in the media inside the pores of the hydrogel, and diffusion coefficient of TMZ in the media outside the hydrogel, respectively. The parameters of the simulations are as follows: **u**= 0, *ϵ* =0.25, D_e,i_ =*ϵ*^4/3^× 10^−10^ =1.57 × 10^−11^ m^2^/s, D_*j*_ = 10^−9^ m^2^/s, *c*_*i*_ =0, *c*_*j*_ = 1 to 1000 μM, *c*_*p,i*_ =0, *μ* = 0.00089 Pa s, **g** =0, and *ρ* =1000 kg/m^3. [53]^ The boundaries of the simulated domain were considered to be impermeable and non-slipping. The computational model was divided into 13,698 triangular meshes to obtain the grid-independent study. Cutlines 1 and 2 were drawn on the simulated domain to analyze the results from the simulation (**Figure S1B**), where cutlines 1 and 2 were used to determine the diffusion of molecules in the hydrogel with respect to incubation time (**Figure S1C-D**) and molecule concentrations (**Figure S1E**). The spatiotemporal distribution of a solution (with a concentration of 1000 μM) was generated to visualize the diffusion of molecules in the hydrogel (**Figure S1F**). The simulation was validated using a hydrogel immersed in a 120 μM Rhodamine solution to evaluate the diffusion dynamics of Rhodamine in the hydrogel (**Figure S1G**).

### Cell Culture

LN229 (ATCC, LN-229 - CRL-2611) and PDM140 (ATCC, HCM-BROD-0195-C71) cells were cultured in neurobasal and DMEM/F12 media (1:1 ratio) containing 2% 1X non-essential amino acid, 2% 1X sodium pyruvate, 2% 1X sodium bicarbonate, 5% 25mM HEPES, 2% 1X glutamax, and 2% mL 1X Antibiotic Antimycotic. U87 (ATCC, HTB-14) cells were cultured in DMEM + glutamax containing 10 % Fetal Bovine Serum (FBS) and 1 % penicillin/streptomycin. Primary human pericytes (ScienceCell, Human Brain Vascular Pericytes) were cultured in pericyte media (ScienceCell) containing 2% FBS (ScienceCell), 1% pericyte growth supplement (ScienceCell), and 1% penicillin/streptomycin solution (ScienceCell). All the cells were cultured according to the manufacturer ‘s recommendation, and, for subculturing, the cells were trypsinized with Accutase (Innovative Cell Technologies, Inc.). For all the experiments, the cells were used before reaching passage number 6.

### Preparation, characterization, and TMZ treatment of GBM, pericytes, and GBM/pericytes bioconstructs

#### (i) Formation and characterization of GBM spheroids

MicroTissues® 3D Petri Dish® micro molds (256-Small Spheroids, MicroTissues Inc.) were used to fabricate agarose microwell arrays in which uniform GBM spheroids were generated. Following the manufacturer ‘s protocol, we fabricated agarose microwell arrays with 256 circular wells (diameter: 300 μm; and height: 800 μm), which were placed in a 12-well tissue culture plate containing 2.5 mL of cell culture medium. Before seeding the cells in the agarose microwell array, the cell culture medium was completely removed, and 190 μL of cell suspension containing 107,000 cells was gently pipetted into the agarose microwell array. Once the cells settled in the microwells after 10 minutes, 2.5 mL of cell culture medium was added to the well of the 12-well tissue culture plate to immerse the agarose microwell array. The 12-well tissue culture plates were incubated at 37°C with 5% CO_2_ and cultured for 3 days by replacing 50% of the cell culture media daily. The spheroids were imaged daily using confocal microscopy, and their viability and morphology (area, aspect ratio, and circularity) were analyzed using the ImageJ software and the FiJi plug-in.

#### (ii) GBM spheroids, pericytes, and GBM/pericytes bioconstructs generation, culture, and TMZ treatment

To generate cylindrical hydrogel bioconstructs (diameter: 5 mm; and height: 5 mm), we cast 98 μL of prepolymer solution containing GBM spheroids (128 spheroids corresponding to 5.3×10^4^ cells per bioconstruct) and/or pericytes (1.07×10^5^ cells per bioconstruct) in the cylindrical shell PMMA mold, and the cell-laden prepolymer solution was crosslinked with UV with a power intensity of 500 mW/cm^2^ for 15 seconds. After crosslinking, the hydrogel bioconstruct were gently removed from the PMMA mold and cultured at 37°C with 5% CO_2_ in 9 mL of media (monoculture: GBM or pericyte media; and co-culture: 1:1 ratio of GBM:pericyte media) in 6-well plates for 14 days, with one complete media change on day 7. On day 7, the GBM or pericytes bioconstructs were treated with 1, 10, 50, 100, 250, 500, and 1000 μM TMZ solutions, and the GBM/pericyte bioconstructs, LN229-pericytes, PDM140-pericytes, and U87-pericytes, were treated with 1000 μM, 100 μM, and 1000 μM TMZ solutions, respectively. TMZ was dissolved in DMSO and then diluted in cell culture media to reach a final DMSO concentration ≤0.1%. The viability of all bioconstructs was evaluated on days 1, 7, and 14. The bioconstructs were imaged on days 1, 7, and 14 using confocal microscopy, and the morphologies of the spheroids (area, aspect ratio, and circularity) were analyzed using the ImageJ software and the FiJi plug-in.

### Cell Viability

The viability of the GBM or pericytes bioconstructs was assessed using the PrestoBlue assay (Invitrogen A13262) following the manufacturer ‘s recommendation with slight modifications. The PrestoBlue reagent was diluted with cell culture media at a final concentration of 10% (v/v). The bioconstructs were placed in a 48-well plate with 300 μL of PrestoBlue solution and incubated for 2 hours at 37°C. After the incubation, 200 μL of the PrestoBlue solution was transferred to a 96-well plate for fluorescence measurement (excitation: 540 nm; emission: 610 nm). The normalized fluorescence intensity (FI) was calculated for each sample by subtracting the FI of blank wells (not containing cells) from the FI of test samples. In addition, the viability of GBM or pericytes bioconstruct was confirmed using confocal image analysis. The bioconstructs were stained using LIVE/DEAD Viability/Cytotoxicity Kit (Invitrogen, cat. # L3224) and Hoechst 33342 solution (Thermo Scientific, cat. # 62249). Briefly, the bioconstructs were incubated for 2 hours at 37°C with 2 μM of Calcein AM, 4 μM Ethidium Homodimer, and 0.1 μg/mL of Hoechst 33342 solution (Thermo Scientific, cat. # 62249). The viability of GBM spheroids or pericytes was assessed using the ImageJ software and the Fiji plug-in. The total number of cells was determined by counting the Hoechst-stained nuclei. The number of live (NL) cells was calculated by counting the cells with green and blue fluorescence. Similarly, the number of dead (ND) cells was calculated by counting the cells with red and blue fluorescence. Finally, the cell viability was calculated with equation (8):

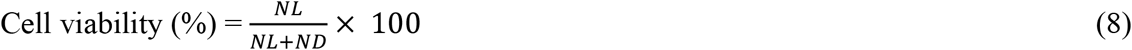

The viability of the GBM/pericytes bioconstructs was assessed using fluorescent confocal image analysis. Before generating the GBM/pericytes bioconstructs, the pericytes were stained with a live cell tracker (CellTracker™ Deep Red Dye, C34565) following the manufacturer ‘s recommendation. Then, the GBM/pericytes bioconstructs were stained using the LIVE/DEAD Viability/Cytotoxicity Kit. The viability of GBM cells and pericytes within the GBM/pericyte bioconstructs was assessed using the ImageJ software and the Fiji plug-in. First, the fluorescence channels (green (live cells), red (dead cells), and magenta (pericytes)) were separated from the raw confocal images and changed to 8-bit greyscale. Using the automated area calculator in the ImageJ software, the area of dead pericytes (DP) was determined by the overlapping areas of all pericytes (magenta) and dead cells (red), and the area of live pericytes (LP) was calculated by subtracting the area of DP from the area of all pericytes (magenta). The area of dead GBM cells (DG) was calculated by subtracting the area of DP from the area of all dead cells (red), and the area of live GBM cells (LG) was calculated by subtracting the area of LP channel from the area of all live cells (green). Lastly, the viabilities of GBM cells and pericytes were calculated with equations (9) and (10), respectively:

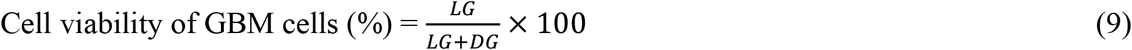

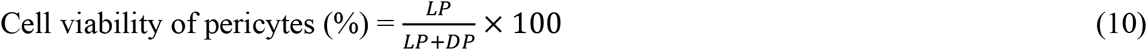

### Spheroid morphology

Spheroid areas, aspect ratios, and circularity were quantified using the ImageJ software and the Fiji plug-in. All spheroid images were changed to 8-bit grayscale. The threshold value was adjusted to establish a clear boundary between the spheroids and the background. The area of a spheroid was measured using the “Analyze Particles” function. To determine the aspect ratio of a spheroid, the “Fit Ellipses” option in the “Analyze Particles” function was used to calculate the major and minor axes of a spheroid, and the aspect ratio was calculated by dividing the major axis by the minor axis. To determine the circularity of a spheroid, the “Analyze Particles” function was used to measure the area and perimeter of a spheroid, and the circularity of a spheroid was calculated with equation (11):

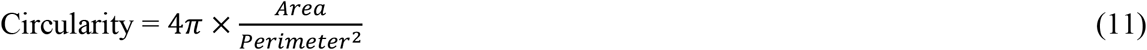

#### qRT-PCR analysis

The mRNA expression of GADD45a, PTEN, MSH6, and CCL5 was quantified for GBM or pericytes bioconstructs on days 1, 7, and 14 of culture using qRT-PCR.

The RNA was extracted from the bioconstruct using the RNeasy Plus Mini Kit (Qiagen, cat # 74134). The QuantiTect Reverse Transcription Kit (Qiagen, cat # 205311) was used to reverse-transcribe the total RNA to complementary DNA (cDNA) with the corresponding primers for target human genes (GADD45a, PTEN, MSH6, and CCL5) and housekeeping genes (GAPDH and β-actin) (primer sequences used are listed in **Table S1**). The reaction conditions for the qRT-PCR included an initial denaturation step at 95°C for 5 s. This was followed by a recurring denaturation step at 95°C for 5 seconds and an amplification step at 60°C for 10 s up to 45 cycles. The QuantStudio 3 software (Thermo Fisher Scientific) was used to compute the Ct values from the melt curves. Gene expression analysis was performed using the 2^-ΔΔCt^ method.^[54]^ The difference between the Ct values of the target genes and the housekeeping genes, or ΔCt, were calculated. The relative mRNA expression levels were normalized by subtracting the ΔCt of the control sample (day 1) from the ΔCt of the experimental sample (day 7 and day 14 with and without TMZ) to obtain ΔΔCt values. These values were used to generate a heat map of the relative mRNA expression changes for GADD45a, PTEN, MSH6, and CCL5.

### Statistical Analysis

All statistical analyses were performed using the GraphPad Prism 9 software. Results are expressed as mean ± standard deviation from triplicate experiments. All data was tested for normality using the Shapiro-Wilk and Shapiro-Francia tests. The area under the curve was calculated using the area under the curve analysis (AUC) in Prism 9 software, where cell viability was plotted in function of TMZ concentration, and a non-linear fit was applied to generate the function used to calculate the AUC. When applicable, analyses for significance were performed using a parametric or non-parametric analysis of variance (ANOVA) with the post-hoc Tukey multiple comparison test. Statistical significance was defined by p<0.05 for two-sided p-values.

## Supporting Information

Supporting Information is available from the Wiley Online Library or from the author.

## Acknowledgements

The authors acknowledge the funding support from the Terasaki Institute for Biomedical Innovation. The authors appreciate the Indian Institute of Technology Guwahati for providing the COMSOL simulations and results.

