## Supplementary file for "Microphysiological system modeling pericyte-induced temozolomide resistance in glioblastoma"

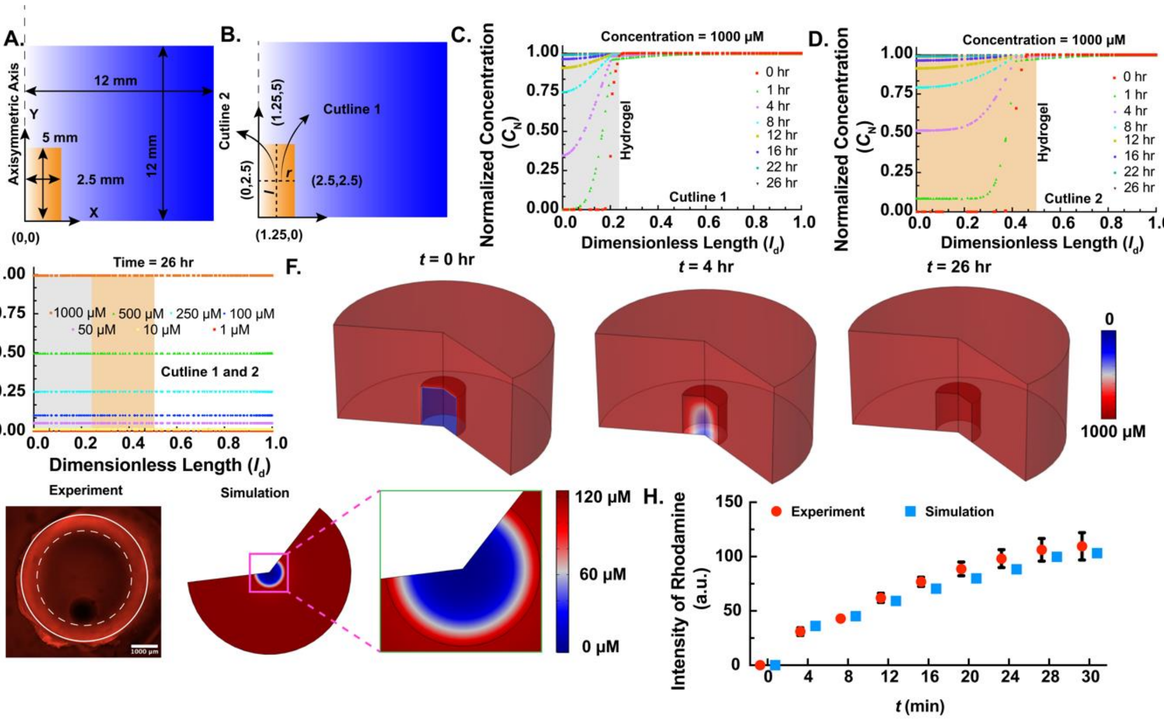


**Figure S1: Validation with computational fluid dynamics (CFD) simulation and parametric study. A.** Dimension of the simulated domain. **B.** Cutline 1 and 2. **C-D.** Distribution of normalized concentration of TMZ with dimensionless length through cutline 1 and 2 at a concentration of TMZ, *C* = 1000 μM, respectively. **E.** Distribution of normalized concentration of TMZ with dimensionless length through cutline 1 and 2 at a concentration of TMZ, *C* = 1000, 500, 250, 100, 500, 10, and 1μM and *t* = 26 hr. **F.** Spatiotemporal distribution of TMZ concentration into a hydrogel at *t* = 0, 4, and 26 hr. **G.** Diffusion of

Rhodamine in the hydrogel with time and corresponding simulation. **H.** Validation of the Rhodamine intensity variation with simulation.

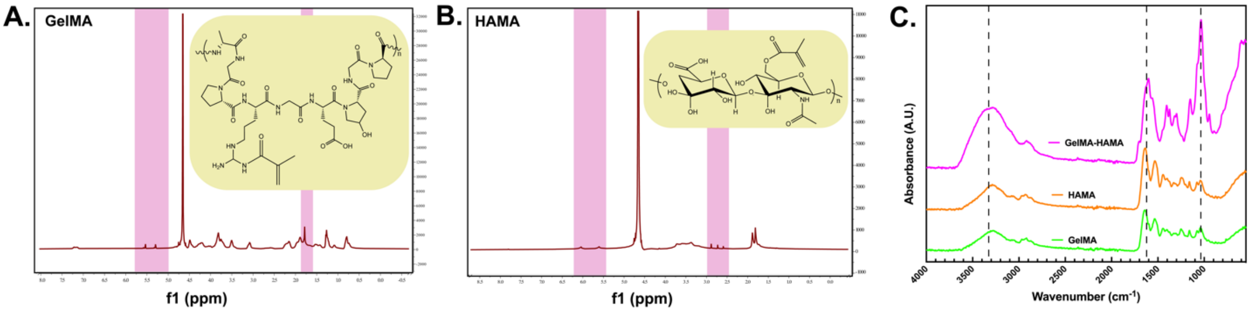


**Figure S2: Chemical characterization of GBM/brain tissue mimicking GelMA-HAMA hydrogel. A.** NMR of GelMA. **B.** NMR of HAMA. **C.** FTIR of GelMA, HAMA, and GelMA-HAMA.

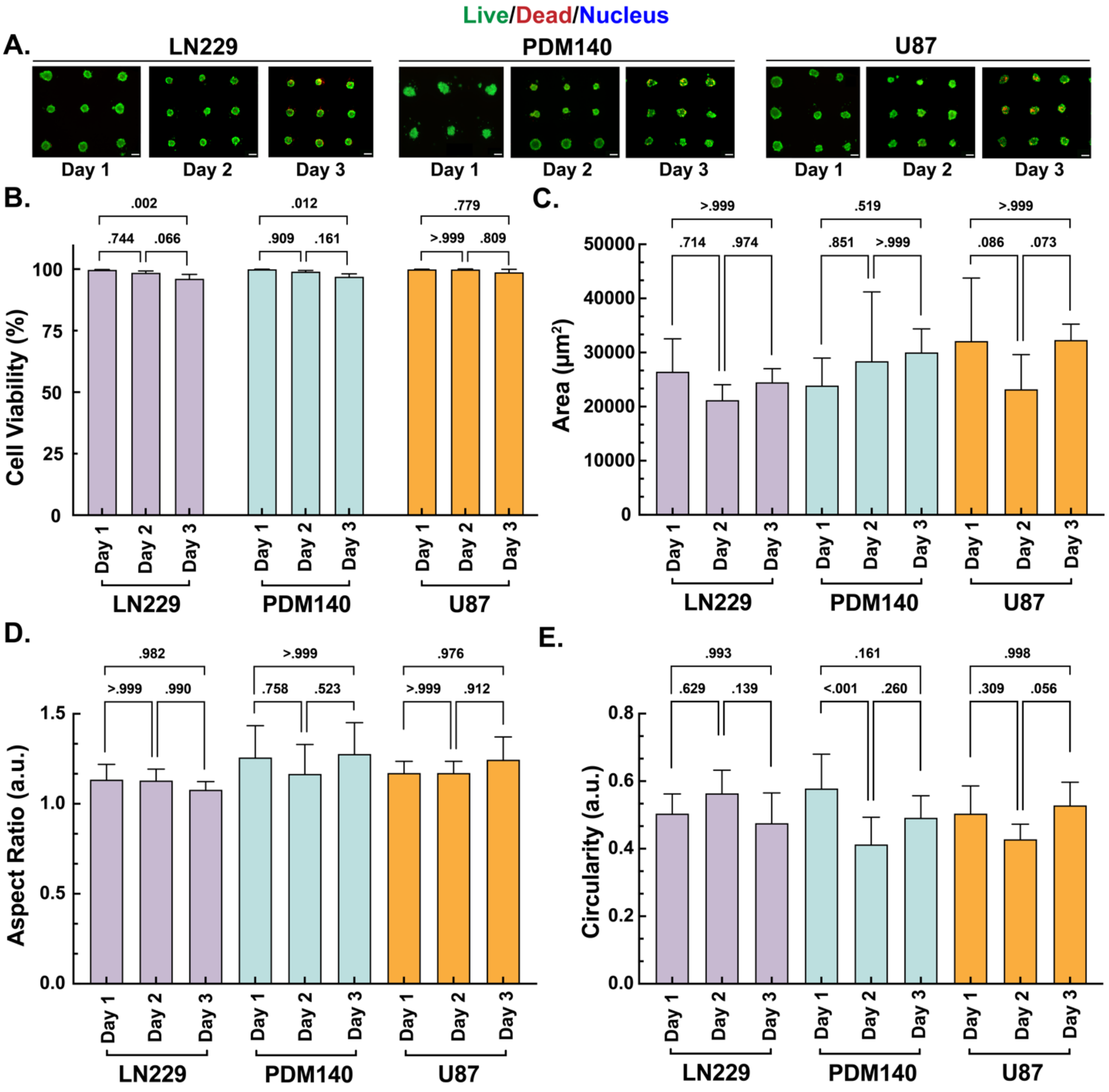


**Figure S3: Formation of GBM spheroids using LN229, PDM140, and U87 cells. A.** Live (green) and dead (red) staining of LN229, PDM140, and U87 spheroid on days 1, 2, and 3 (scale bar = 150 μm). **B.** Viability of LN229, PDM140, and U87 spheroids on days 1, 2, and 3. **C.** Area of LN229, PDM140, and U87 spheroids on days 1, 2, and 3. **D.** Aspect ratio of LN229, PDM140, and U87 spheroids on days 1, 2, and 3. **E.** Circularity of LN229, PDM140, and U87 spheroids on days 1, 2, and 3. Statistical test used with * $p < 0.05$, ** $p < 0.01$, *** $p < 0.001$.

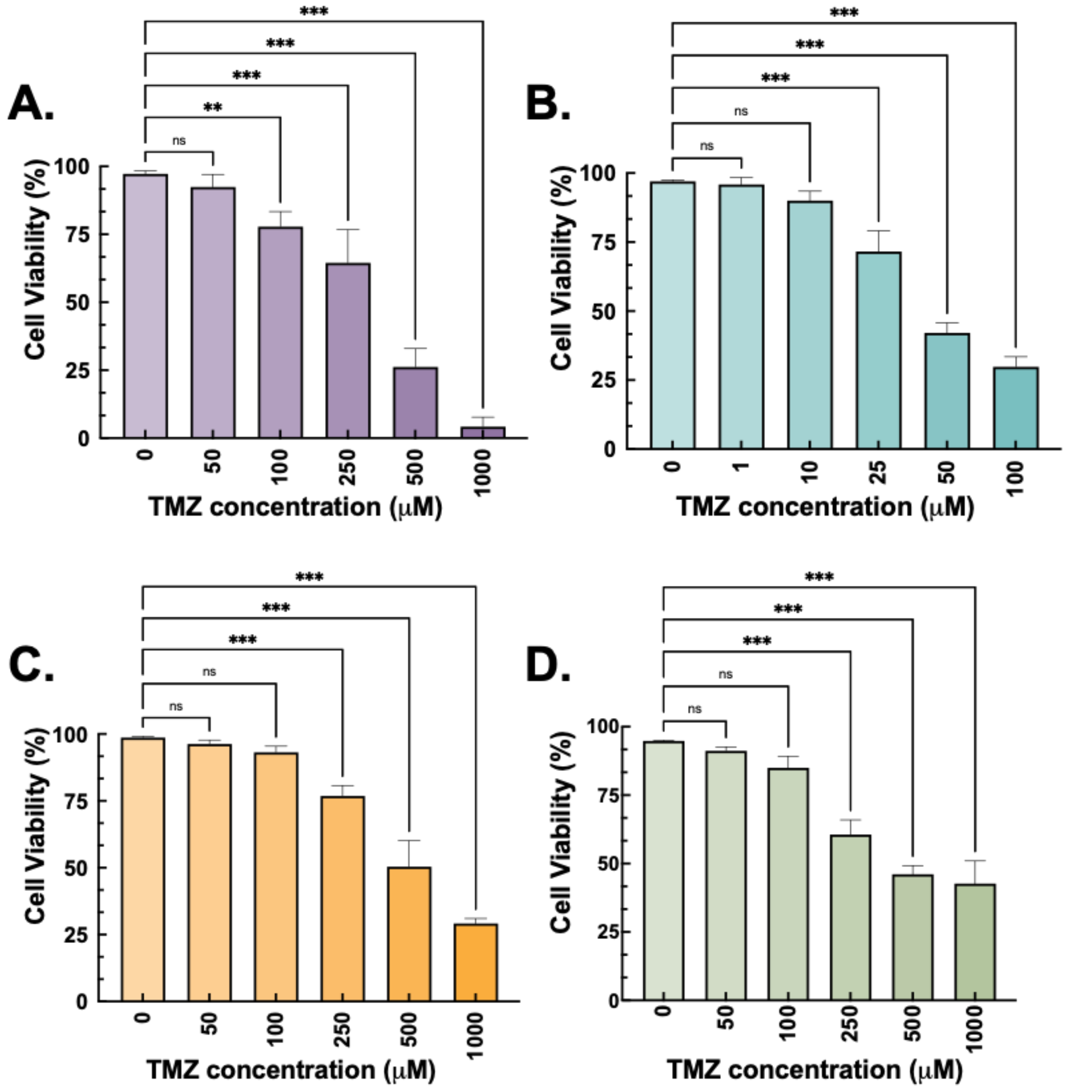


**Figure S4:** Confocal image analysis of cell viability for **A.** LN229, **B.** PDM140, **C.** U87, and **D.** pericytes treated with different concentrations of TMZ.

**Table S1:** Primer sequences used for qRT-PCR

| Gene | | Sequences |
|---|---|---|
| GADD45a | Forward | CTG GAG GAA GTG CTC AGC AAA G |
| | Reverse | AGA GCC ACA TCT CTG TCG TCG T |
| PTEN | Forward | TGA GTT CCC TCA GCC GTT ACC T |
| | Reverse | GAG GTT TCC TCT GGT CCT GGT A |
| HSH6 | Forward | AAG GAC TGG CAG TCT GCT GTA G |
| | Reverse | CGG CAA CAG AAT TAC TGG GCG A |
| CCL5 | Forward | CCT GCT GCT TTG CCT ACA TTG C |
| | Reverse | ACA CAC TTG GCG GTT CTT TCG G |
| GAPDH | Forward | GTC TCC TCT GAC TTC AAC AGC G |
| | Reverse | ACC ACC CTG TTG CTG TAG CCA A |
| β-actin | Forward | CAC CAT TGG CAA TGA GCG GTT C |
| | Reverse | AGG TCT TTG CGG ATG TCC ACG T |
